# Cryo-EM Structure of HRSL Domain Reveals Activating Crossed Helices at the Core of GCN2

**DOI:** 10.1101/2024.04.24.591037

**Authors:** Kristina Solorio-Kirpichyan, Xiao Fan, Dmitrij Golovenko, Andrei A. Korostelev, Nieng Yan, Alexei Korennykh

## Abstract

GCN2 is a conserved receptor kinase activating the Integrated Stress Response (ISR) in eukaryotic cells. The ISR kinases detect accumulation of stress molecules and reprogram translation from basal tasks to preferred production of cytoprotective proteins. GCN2 stands out evolutionarily among all protein kinases due to the presence of a <u>h</u>istidyl t<u>R</u>NA <u>s</u>ynthetase-like (HRSL) domain, which arises only in GCN2 and is located next to the kinase domain. How HRSL contributes to GCN2 signaling remains unknown. Here we report a 3.2 Å cryo-EM structure of HRSL from thermotolerant yeast *Kluyveromyces marxianus*. This structure shows a constitutive symmetrical homodimer featuring a compact helical-bundle structure at the junction between HRSL and kinase domains, in the core of the receptor. Mutagenesis demonstrates that this junction structure activates GCN2 and indicates that our cryo-EM structure captures the active signaling state of HRSL. Based on these results, we put forward a GCN2 regulation mechanism, where HRSL drives the formation of activated kinase dimers. Remaining domains of GCN2 have the opposite role and in the absence of stress they help keep GCN2 basally inactive. This autoinhibitory activity is relieved upon stress ligand binding. We propose that the opposing action of HRSL and additional GCN2 domains thus yields a regulated ISR receptor.

**Significance statement:** Regulation of protein synthesis (translation) is a central mechanism by which eukaryotic cells adapt to stressful conditions. In starving cells, this translational adaptation is achieved via the receptor kinase GCN2, which stays inactive under normal conditions, but is switched on under stress. The molecular mechanism of GCN2 switching is not well understood due to the presence of a structurally and biochemically uncharacterized <u>h</u>istidyl t<u>R</u>NA <u>s</u>ynthetase-like domain (HRSL) at the core of GCN2. Here we use single-particle cryo-EM and biochemistry to elucidate the structure and function of HRSL. We identify a structure at the kinase/HRSL interface, which forms crossed helices and helps position GCN2 kinase domains for activation. These data clarify the molecular mechanism of GCN2 regulation.

## Introduction

GCN2 is a stress kinase from the family of four translation initiation factor 2α (eIF2α) kinases (1, 2). These kinases sense signaling molecules produced specifically in stressed cells and phosphorylate a regulatory serine residue of translation initiation factor eIF2α. Phosphorylation of eIF2α inhibits the initiation step of protein synthesis and activates the Integrated Stress Response (ISR) to allow cell adaptation to stress. The ISR attempts to re-establish normal cell function by reprogramming gene expression, and if homeostasis cannot be achieved, the ISR activates apoptosis (2).

The four ISR receptor kinases are GCN2, HRI, PERK and PKR. They contain homologous kinase domains, which recognize and phosphorylate eIF2α. The kinase domains are linked to different stress-sensing and regulatory domains for each family member. These domains allow GCN2 to sense the nutrient status of a cell (2, 3), HRI to sense heme (4), PERK to sense unfolded proteins (5), and PKR to sense harmful double-stranded RNA (6, 7). GCN2^-/-^ mice have few pups under low-leucine diet and develop severe liver steatosis when under amino acid deprivation due to importance of GCN2 in metabolic regulation of nutrients (8, 9). Depending on the type of training, GCN2^-/-^ mice can exhibit enhanced or impaired learning ability, supporting the broad role of the ISR in cognitive processes (2, 10, 11).

Structural and biochemical studies have shown that regulation of the ISR kinases uses homo-dimerization, although the precise mechanisms of dimerization coupling to the ligand-sensing domains remain poorly understood for all ISR kinases (6). Structures of every major domain are available for two of the four receptors: PERK and PKR. These structures with biochemical and other studies provide perhaps the most complete models of regulation among ISR kinases (5–7, 12–14). GCN2 is the largest ISR receptor kinase featuring five globular domains. Structures of three GCN2 domains have been reported. At the N-terminus, GCN2 contains the <u>R</u>ing finger-<u>W</u>D repeat-DEAD-like helicase <u>D</u>omain (RWD), which forms a stable homodimer and binds to GCN2 partner proteins GCN1/GCN20 (15–17) (Fig. 1B, PDB ID 7E2M). At the C-terminus, GCN2 contains the <u>C</u>-<u>T</u>erminal <u>D</u>omain (CTD), which also forms a homodimer and apparently has two functions: binding to the ribosome and autoinhibition of the kinase (18, 19) (Fig 1B, PDB 4OTM). Finally, structures of the kinase domain have been determined for two functionally relevant dimeric states. They revealed a head-to-head inactive dimer (20) (Fig 1B. PDB 1ZYD) and a back-to-back active dimer, which is similar to active dimers formed by other protein kinases (21, 22) (Fig 1B, PDB 6N3O and 7QQ6). Structure of GCN2 <u>p</u>seudo-<u>k</u>inase <u>d</u>omain (ΨKD), proposed to help GCN2 activation (23), remains unknown. The largest GCN2 domain, sharing 22% sequence identity with histidyl tRNA synthetase HisRS (<u>H</u>is<u>RS</u>-<u>L</u>ike domain, HRSL; Fig. 1A, B), also remains structurally uncharacterized.

**Figure 1.**
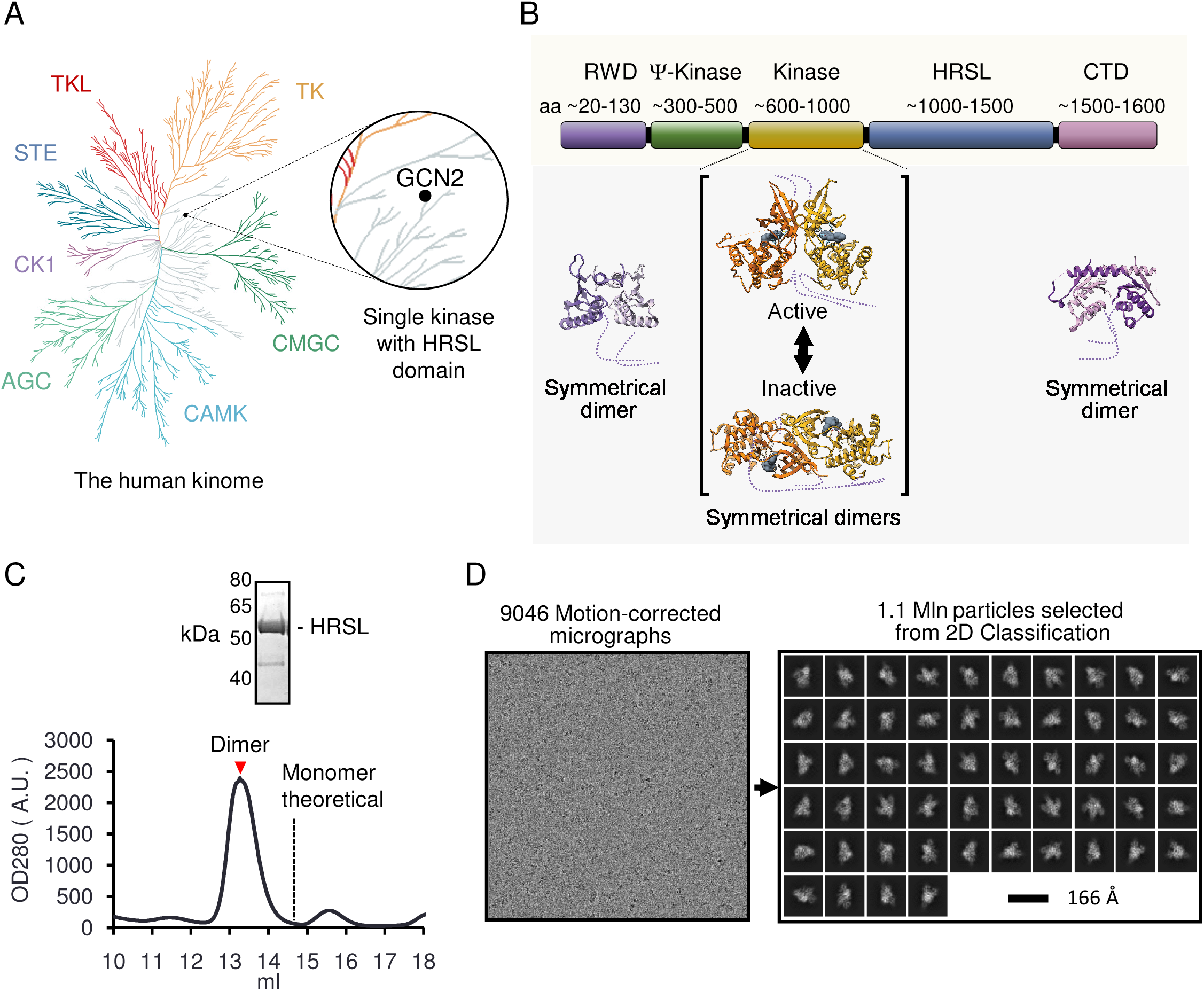
Phylogenetic attributes of GCN2 and characterization of HRSL domain. A) Location of GCN2 within human kinome. B) Domain architecture of GCN2 with available crystal structures superimposed. PDB codes of shown structure: RWD 7E2M; kinase 6N3O (active dimer) 1ZYD (inactive dimer); CTD 4OTM. C) Left: Size exclusion chromatography FPLC trace of purified GCN2 HRSL. GCN2 HRSL runs at a molecular weight consistent with a dimer at ∼120 kDa. Right: Coomassie stained SDS-PAGE with purified GCN2 HRSL collected from indicated fraction (red arrow). D) Left: Representative motion-corrected cryo-EM micrograph for GCN2 HRSL. Right: Classes selected from final round of 2D classification used for initial 3D reconstruction.

The deficit of structural information about major domains of GCN2 and the absence of tRNA synthetase-like domains in other, better-studied protein kinases (Fig. 1A) (24) limits our understanding of the molecular mechanism of GCN2. To lift this limitation, here we present single-particle cryo-EM and biochemical analyses of GCN2 HRSL, offering insights into the function of HRSL.

## Results

### Cryo-EM Structure of GCN2 HRSL Domain Reveals an Intertwined Homodimer

To characterize the HRSL domain, we cloned GCN2 from three species—human, *S. cerevisiae*, and thermotolerant yeast *Kluyveromyces marxianus*—and subsequently purified the corresponding HRSL domains, both individually and with flanking domains of various length. Only *Kluyveromyces* sequence yielded a soluble homogeneous HRSL domain suitable for structural studies (residues 996-1511; Methods). HRSL migrated as a stable homodimer (Fig. 1C). Milligram quantities of *Kluyveromyces* HRSL were produced and tested in high-throughput crystallization screens, without success. By contrast, well-dispersed Quantifoil Cu grid samples suitable for cryo-EM analysis were successfully prepared (Fig. 1D).

We determined a 3.2 Å cryo-EM structure of GCN2 HRSL (Fig. S1; Table S1), in which most side chains are well resolved (Fig. 2A-B). To build the structure of GCN2 HRSL, we generated an initial AlphaFold model from protein sequence (25) (Methods). The predicted model, containing a crossed homodimer formed by intertwined monomers, had a nearly perfect fit into the map and required only minor corrections of several side chains and a conservative all-atom refinement to achieve the final structure (Fig. 2A-D; Methods). Compared to histidyl tRNA synthetase HisRS (26), GCN2 HRSL has an additional N-terminal structure comprised of bundled alpha-helices (Fig. 2C-D, boxed). Truncation of these N-terminal helices in a mutant Δ(V996-N1009) caused a loss of the N-terminal density, providing experimental verification of correct helix placement in our model (Fig. 2E; Fig. S2 A-C; Table S2).

**Figure 2.**
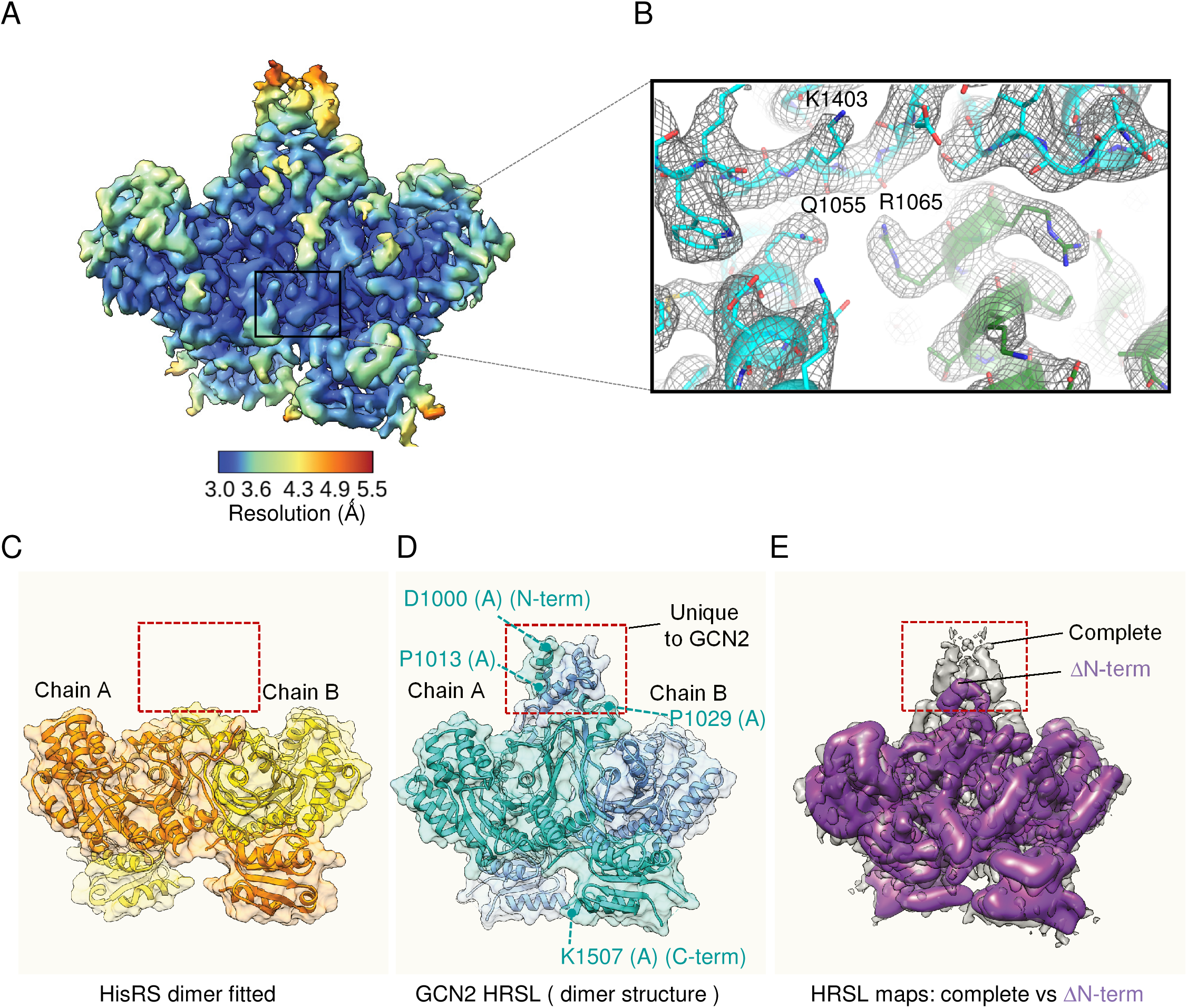
Cryo-EM structure analysis of GCN2 HRSL domain. A) Final sharpened GCN2 HRSL cryo-EM map generated from CryoSPARC, surface colored by local resolution based on gold standard FSC threshold 0.143. B) Representative densities of amino acid backbone and side chains. C) Structure tRNA synthetase HisRS (PDB ID 4RDX). D) Structure of GCN2 HRSL. Boxed region shows a unique subdomain absent in HisRS. E) Cryo-EM map for ΔN-terminal GCN2 HRSL dimer (purple) superimposed onto the cryo-EM map of GCN2 HRSL (grey), Tables S1 and S2.

### Conservation Analysis of HRSL Surface Electrostatics Suggests a Loss of tRNA Complementarity

Based on sequence homology with His-tRNA synthetase HisRS, HRSL domain of GCN2 was initially proposed to also bind free tRNAs, modestly preferring deacylated tRNAs, to sense the nutritional status of a cell (27). An alternative and perhaps complementary recent model proposes that GCN2 binds the P-stalks of stalled ribosomes, using the ribosome to sense whether the cell undergoes a translational stress (28, 29). The HRSL domain is linked directly to the kinase domain and thus it should play a role in regulation. Indeed, strong activation of GCN2 kinase by ribosomes depends on HRSL (29).

Here we took advantage of our structure to re-evaluate the tRNA-sensing model. We analyzed and compared the tRNA-interaction site of HisRS with equivalent surface of GCN2 HRSL. Considering that RNA-binding surfaces are usually positively charged to recognize negatively charged phosphates of RNA backbone, we computed conserved electrostatic potentials (Methods). Our analysis takes into account 3D structures and evolutionary conservation of amino acid charges obtained from multiple sequence alignments of more than 190 eukaryotes, from yeasts to human. HisRS exhibits an evolutionarily conserved positive surface potential that tracks well with tRNA (Fig. 3A-B). By contrast, HRSL of GCN2 lacks global electrostatic complementarity with tRNA and harbors a conserved antagonizing negatively charged patch at the putative binding site for tRNA anticodon stem-loop (ASL) (Fig. 3C-D).

**Figure 3.**
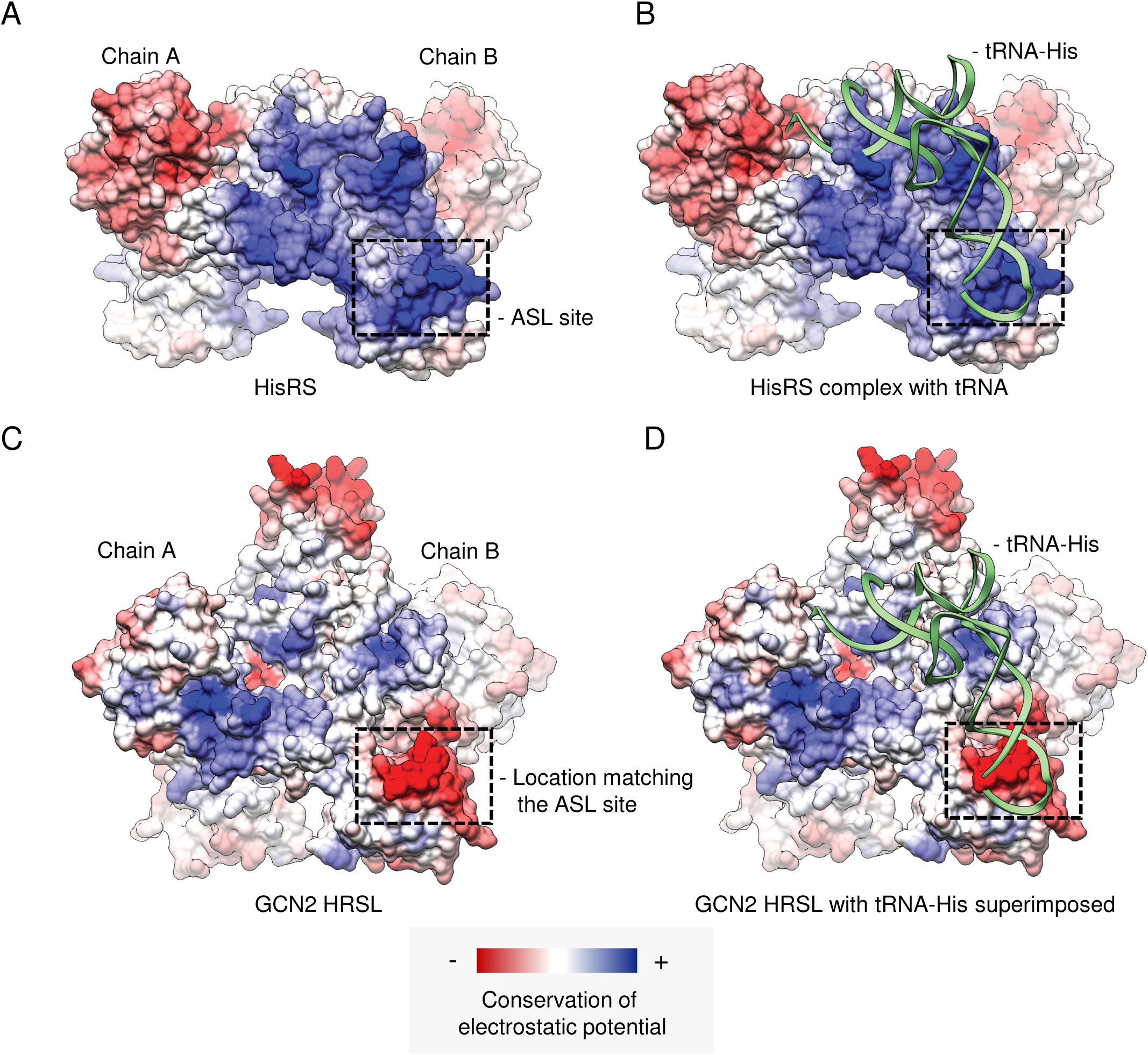
Comparison of surface electrostatics conservation in tRNA synthetase HisRS versus GCN2 HRSL. A) HisRS structure (PDB: 4RDX) colored by conservation of electrostatic potential (Methods). Dotted box indicates <u>a</u>nticodon <u>s</u>tem loop (ASL) site. Bound tRNA is hidden for visualization of surface charge. B) Structure from (A) with bound tRNA (green). C) GCN2 HRSL structure colored by conservation of electrostatic potential. D) Structure from (C) with tRNA (green) docked to the surface based on structure alignment with HisRS (B).

### HRSL Connects to the Kinase Domain via Dimerized <u>J</u>unction α-<u>H</u>elices (JH)

The N-terminal part of HRSL (aa 1000-1029) contains two sequential α-helices, which are absent in tRNA synthetases and form a symmetrical parallel homodimer (bundle) in the dimeric HRSL structure (Fig. 2B-D; 4A). The bundle is kept by extensive hydrophobic interactions. Helix 1 of one HRSL copy (D1000-P1013, Fig. 2D) is followed by helix 2 (S1014-P1029) placed at an angle. Two symmetry-related helices from the partner chain complete the bundle. The core is held by I1003, L1007, I1010 (helix 1) and W1017, V1021 and L1025 (helix 2) from both HRSL monomers. The first helix has been captured in the previous crystal structure of GCN2 kinase domain, however its role could not be assigned: neither does it pack compactly with the rest of the kinase nor is it located proximally to known functional sites of kinase (20) (Fig. 4B). Our work now shows that the α-helix is a part of HRSL domain, serving as kinase/HRSL <u>J</u>unction <u>H</u>elix (JH).

**Figure 4.**
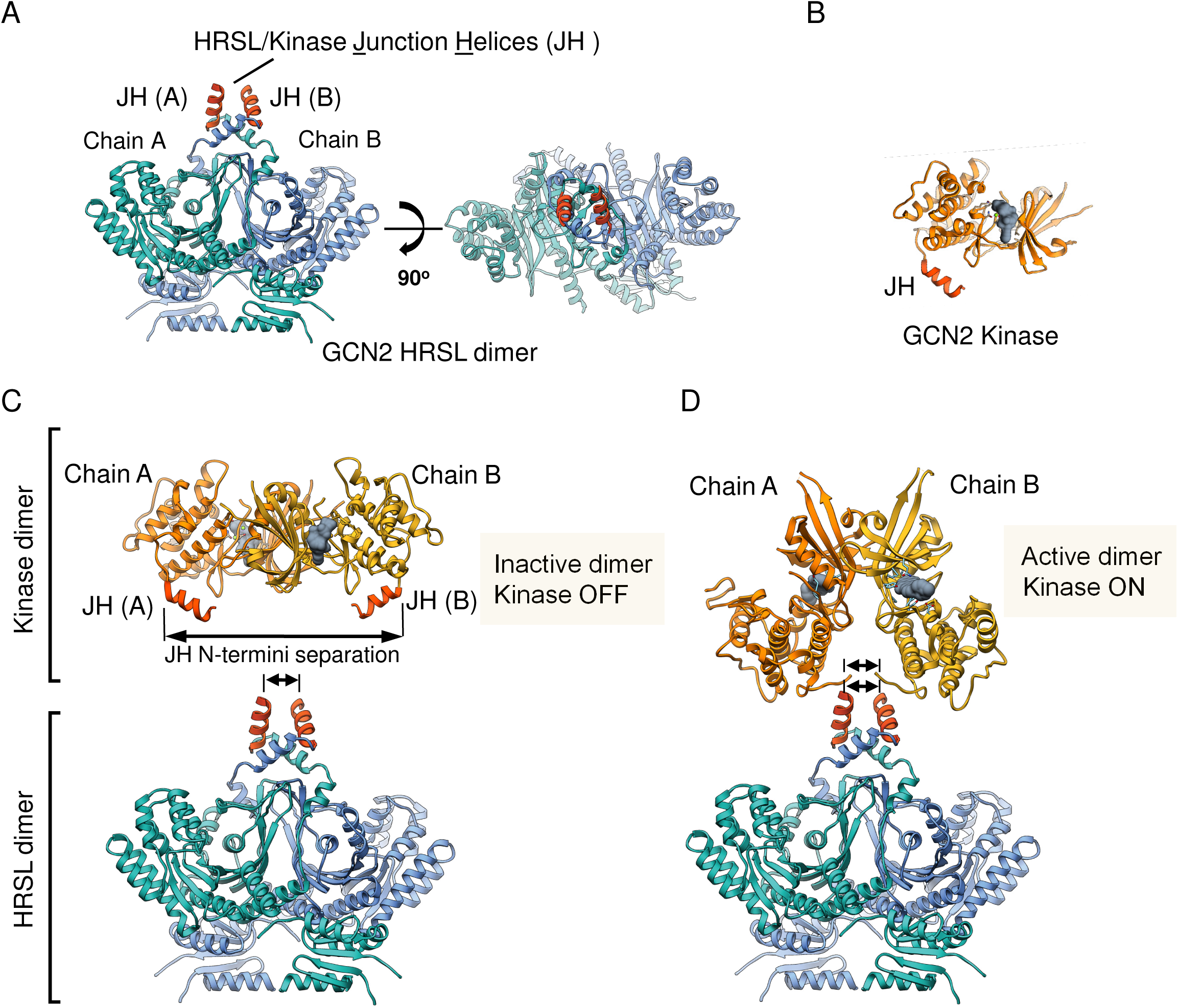
Dimerized α-Helices JH connect HRSL and kinase domains. A) GCN2 HRSL dimer with <u>J</u>unction <u>H</u>elices (JH) colored red for each chain. B) GCN2 kinase domain (monomer-PDB: 1ZYD) with JH colored red. C-D) Structure of inactive (PDB: 1ZYD) and active (PDB: 6N3O) dimers of GCN2 kinase domain next to the dimer of GCN2 HRSL. Distances between equivalent amino acids in kinase and HRSL domains are marked with arrows. The inactive dimer has the kinase domains bound N-lobe-to-N-lobe, with anchor points for JH helices well-separated. The kinase domains adapt the inactive (α-C helix-out) conformation. The active dimer has the kinase domains bound back-to-back, with anchor point for JH helices positioned proximally. The kinase domains adapt the active (α-C helix-in) conformation.

Analogous to the HRSL domain, which exists as a stable homodimer, the kinase domain of GCN2 also forms a homodimer. Structural studies identified two types of dimers for GCN2 kinase. The first type contains the catalytically inactive “OFF” conformation of kinase (PDB ID 1ZYD) (20), whereas the second type contains the catalytically active “ON” conformation of kinase (PDB ID 6N3O) (30). To understand how HRSL could interact with these dimers, we placed either dimeric kinase next to the dimeric HRSL from our cryo-EM structure. The dimer of GCN2 kinases in the “OFF” state is sterically incompatible with our structure due to the large distance between the N-termini of JH in kinase compared to that in HRSL dimer (Fig. 4C). In contrast, structure of the dimeric kinase in “ON” conformation matched well with the HRSL dimer (Fig. 4C-D). The cryo-EM structure of HRSL therefore likely represents the active signaling state of GCN2.

### JH Dimerization Activates GCN2

Dimerization of JH helices at the core of GCN2 suggests their role in regulating the receptor, whereas the steric match with the active kinase dimer (Fig. 4) indicates that JH/JH interaction activates (rather than inhibits) the kinase output. To test this hypothesis, we developed an assay for quantitative biochemical analysis of GCN2 kinase activity based on a previous publication (31). We measured phosphorylation of eIF2α by the GCN2 construct comprising the N-terminally FLAG-tagged kinase/HRSL domains of thermotolerant yeast (residues 578-1511; Table S2) purified from HEK293 cells (Methods). As a control for non-specific interactions with anti-flag resin, we used a mock pulldown from naïve HEK293 cells.

Time-dependent phosphorylation of eIF2α was observed in the presence of GCN2 kinase/HRSL (Fig. S3 A-C; Table S3). Eluates from naïve pulldowns did not cause eIF2α phosphorylation, indicating that the observed kinase activity arises specifically from the transfected GCN2 construct rather than from a stray cellular kinase activity (Fig. S3). To test whether GCN2 kinase/HRSL construct is truly active autonomously or might depend on an accidentally co-purified cellular RNA, we treated GCN2 kinase/HRSL with an excessive (300 nM) concentration of RNase A. RNase A treatment did not inhibit the eFI2α phosphorylation activity of GCN2 kinase/HRSL (Fig. S4). These results with GCN2 kinase/HRSL from thermotolerant yeast agree with those obtained using GCN2 kinase/HRSL from human, which also demonstrated constitutive catalytic activity in the absence of tRNAs and ribosomes (31).

We next tested whether disruption of the JH/JH helical homodimer affects the catalytic activity of GCN2. We tested mutations of hydrophobic residues L1007A, V1021N, and V1021A located at the core of the N-terminal helical bundle in our cryo-EM structure (Fig. 5). As a positive control for activity readout with mutants, we tested a known catalytically dead mutant D848N (31), in which the nucleophilic aspartate required for proton transfer is replaced by non-catalytic isosteric asparagine (Fig. 6A-B; S5). The mutation L1007A was detrimental to activity of GCN2 kinase/HRSL, indicating that the amino acid L1007 forms a functional interaction that helps maintain GCN2 kinases in the active dimeric state. The mutation V1021A was neutral, indicating that although alanine is smaller than valine, it is sufficiently hydrophobic to preserve JH/JH interaction. In contrast, a more invasive mutation V1021N interfered sufficiently to disrupt the kinase activity (Fig. 6A-B; Fig. S5).

**Figure 5.**
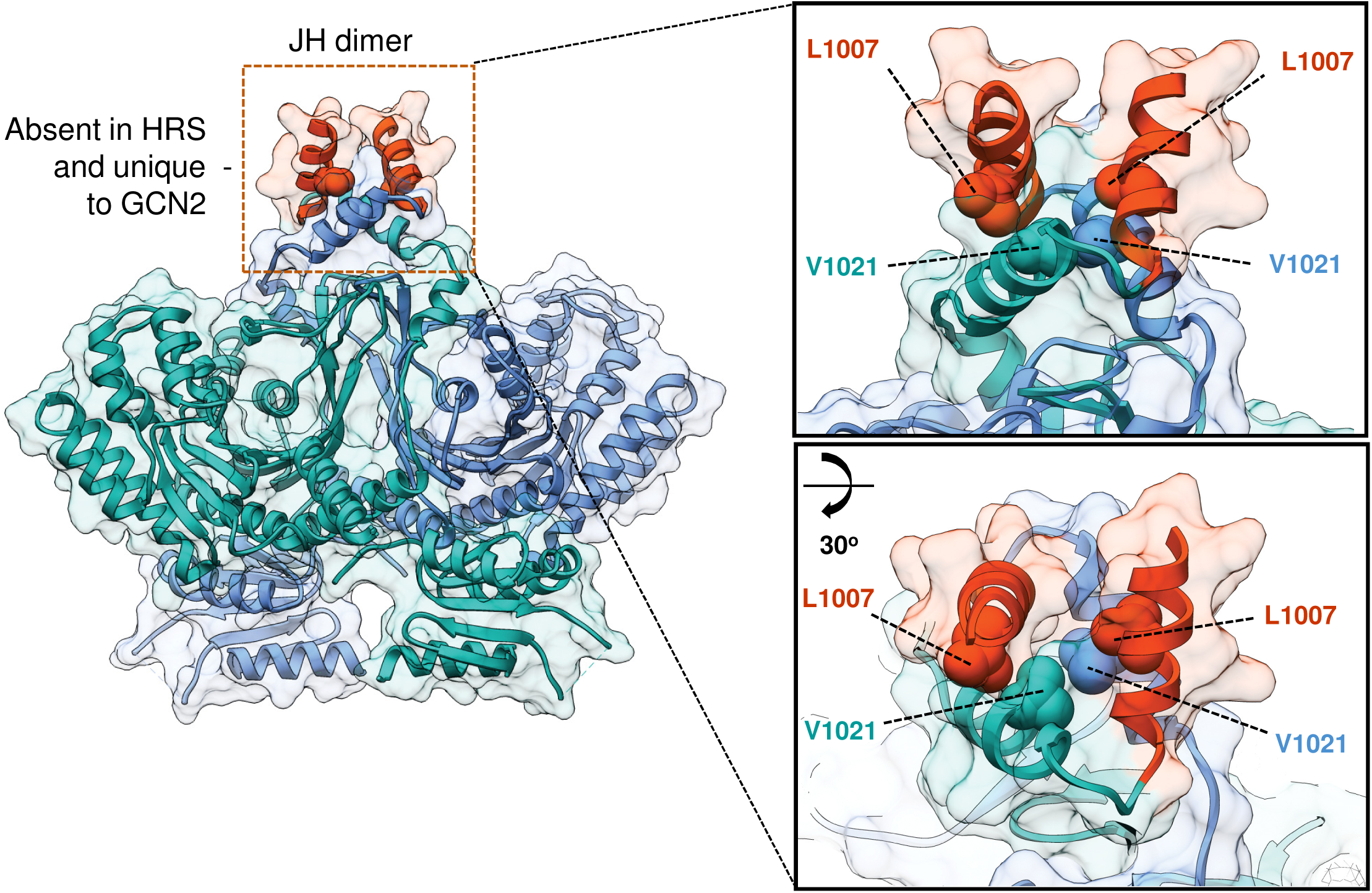
Close-up view of JH/JH interface in GCN2 HRSL domain. Structure of HRSL with JH helices colored orange. The interface between dimerized helices JH is created by several closely packed hydrophobic side chains. We selected hydrophobic amino acid L1007 in JH, and hydrophobic amino acid V1021 located in α-helix immediately downstream of JH for mutagenesis. The α-helix harboring V1021 together with JH contributes to the formation of the α-helical bundle present in GCN2 and absent in HisRS. Two rotationally related close-up views of the bundle structure are showed in boxed panels.

**Figure 6.**
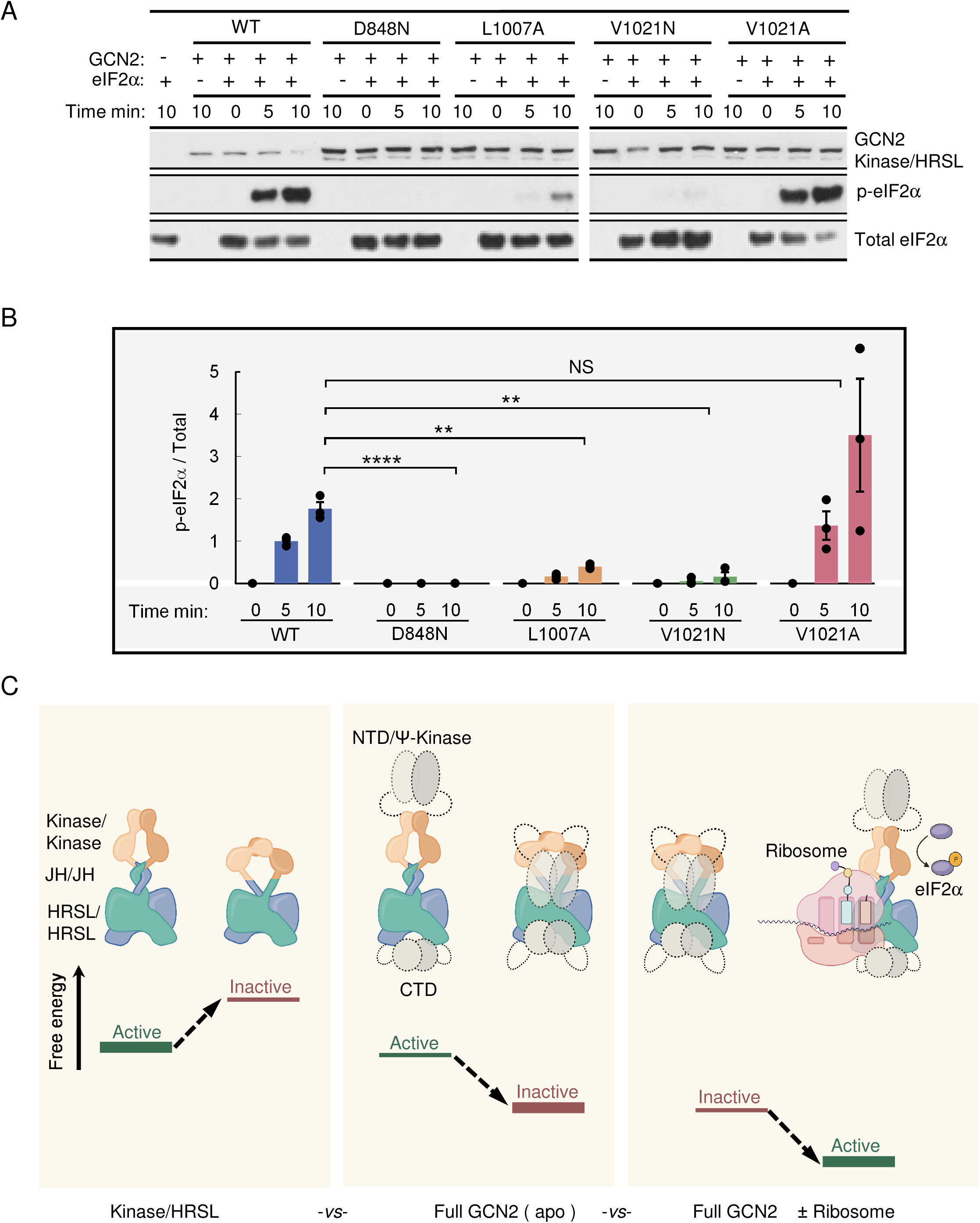
Functional analysis of GCN2 activation by JH and a proposed model of GCN2 regulation. A) In vitro kinase activity for WT and interface mutants (L1007A, V1021A, and V1021N) in GCN2 kinase/HRSL (“K/HRSL”) visualized by western blot. B) Quantification of biochemical data for the mutants D848N, L1007A, V1021A, and V1021N. Statistical significance (P) was calculated using Welch’s two-tailed unpaired t-test: P-value ≤0.05*, ≤0.01**, ≤0.001***, ≤0.0001****, N.S: non-significant. C) Proposed model GCN2 regulation. Relative free energies for kinase/HRSL (left panel), full-length GCN2 (middle panel) and GCN2 bound to the ribosome (right panel) are marked.

## Discussion

Two features stand out in the structure of GCN2 HRSL domain: the loss of electrostatic complementarity to tRNA and the presence of a compact helical bundle formed via dimerization of α-helices JH connecting kinase and HRSL. The loss of the electrostatic complementarity provides a biophysical argument against binding of free tRNAs to GCN2 HRSL as a key stress-sensing step and indicates that sensing stressed ribosomes via ribosomal P-stalk could be the primary mechanism (28, 29). This conclusion is further consistent with the low (∼2 μM) reported K_d_ of tRNA to GCN2 (32) and inability of comparable (5 μM) tRNA concentration to activate GCN2, whereas nanomolar ribosome concentrations are sufficient for GCN2 activation (28, 29). Additionally, a third, unified model of GCN2 activation by both P-stalk and ribosome-bound tRNA cannot be excluded and remains in consideration. In this model, the loss of electrostatic complementarity to tRNA could be a mechanism to desensitize GCN2 to free tRNAs and render the receptor selective toward ribosome-bound tRNAs.

The second major feature of HRSL, JH helix, plays a central role in regulation of GCN2. Dimeric HRSL with dimerized JH/JH region is structurally complementary only to the active configuration of dimeric GCN2 kinase (Fig. 4C vs 4D). This conclusion is supported by biochemical analyses (Fig. 6A-B; Fig. S5) and confirms that our cryo-EM structure represents the kinase-activating state of HRSL. Based on these observations, we propose a mechanism of GCN2 regulation, which predicts free energy relationships and roles of multiple domains of this receptor. The thermodynamic role of HRSL is to favor the catalytically active dimers of GCN2 kinase (Fig. 6C, first panel). If these additional domains were absent, GCN2 would exhibit uncontrollable, persistent activity with harmful consequences, such as apoptosis due to global translational arrest. Therefore, the free energy role of at least some of the additional domains of GCN2, present besides kinase and HRSL, is to counteract the action of HRSL and auto-inhibit the receptor under basal conditions (Fig. 6C, middle panel). The previously proposed auto-inhibitory action of CTD is in line with this model (18, 19). The auto-inhibited state must have JH/JH interface disrupted to sterically match HRSL with the inactive dimer of GCN2 kinase (Fig. 4C) and to account for the deleterious effect of the L1007A and V1021N mutations (Fig. 6A-B). GCN2 becomes activated once it encounters translationally compromised ribosomes. These ribosomes likely outcompete (distract) the auto-inhibiting domains of GCN2, allowing stored free energy in separated JH helices to be released via JH/JH dimerization. The resulting JH/JH dimerization switches GCN2 kinase from the inactive kinase dimer to the active dimer (Fig. 4C,D; Fig. 6C). We predict that the main HRSL domain remains constitutively dimeric in both, active and inactive states of GCN2. This prediction is based on constitutive dimeric state of known tRNA synthetases, and on our experimental observation that HRSL remains dimeric even when the dimerizing JH helices are truncated (Fig. 2E; Fig. S2).

## Database depositions

Coordinates and cryo-EM map were deposited to the PDB and EMDB databases under accession numbers PDB ID 9BF3 and EMD-44489.

## Acknowledgments

We thank Prof. Frederick Hughson and Dr. Sarah Port for providing genomic DNA for *Kluyveromyces marxianus* and for sharing purified SUMO-protease. We are grateful for Prof. Yibin Kang Laboratory for providing HEK293T cells. We would like to acknowledge Princeton’s Imaging and Analysis Center (IAC), which is partially supported through the Princeton Center for Complex Materials (PCCM), and by National Science Foundation (NSF)-MRSEC program (DMR-2011750). We thank Dr. Paul Shao and staff from IAC for their expertise and assistance with cryo-EM data collection. We thank Dr. Matthew Cahn from Princeton University Research Computing for cryo-EM computational support. Molecular graphics and analyses performed with UCSF Chimera, developed by the Resource for Biocomputing, Visualization, and Informatics at the University of California, San Francisco, with support from NIH P41-GM103311.

## Author contributions

K.S. performed cloning, purification, electron microscopy data collection, 6.1 Å data processing for ΔN GCN2 HRSL, initial data processing for WT GCN2 HRSL, and biochemical analyses. X.F. and N.Y. guided electron microscopy sample preparation, electron microscopy data collection, and data processing. D.G. and A.A.K. generated AlphaFold model of GCN2 HRSL from *Kluyveromyces marxianus*, and performed 3.2 Å cryo-EM data processing and structure refinement. K.S. and A.V.K. wrote the manuscript. All authors revised the manuscript and K.S., A.A.K. and A.K. finalized the manuscript.

## Methods

*Kluyveromyces marxianus* GCN2 HRSL sequence (Table S2) was inserted into pETM11-SUMO vector and expressed in *Escherichia coli* and purified to homogeneity by size exclusion fast protein liquid chromatography (FPLC). Cryo-EM samples were prepared with Quantifoil 1.2/1.3 on Cu 300 mesh Holey Carbon film using FEI vitrification robot. Cryo-EM data were collected using Titan Krios G3 cryo Transmission Electron Microscope at 300 kV. Data processing was done with cryoSPARC v4.4.1 (33) (for the 3.2 Å structure) and RELION (34) (for the 6 Å dataset). AlphaFold prediction of initial model of GCN2 HRSL was obtained using a local installation of AlphaFold (25). The model was real-space refined in Phenix and Coot (35–37). In vitro kinase assays were conducted at 32 °C using eIF2α from LifeSpan Biosciences. *Kluyveromyces marxianus* kinase/HRSL domain sequence (Table S2) was inserted into pcDNA4.TO and expressed in HEK293T cells. Cells were grown in DMEM media + 10% FBS (Gibco). eIF2α phosphorylation was measured via western blot using primary antibodies for eIF2α phospho-S51 (Abcam), and total eIF2α (Cell Signaling Technologies). Detailed Methods are provided in SI.

## Supplementary Figure Legends

**Figure S1.**
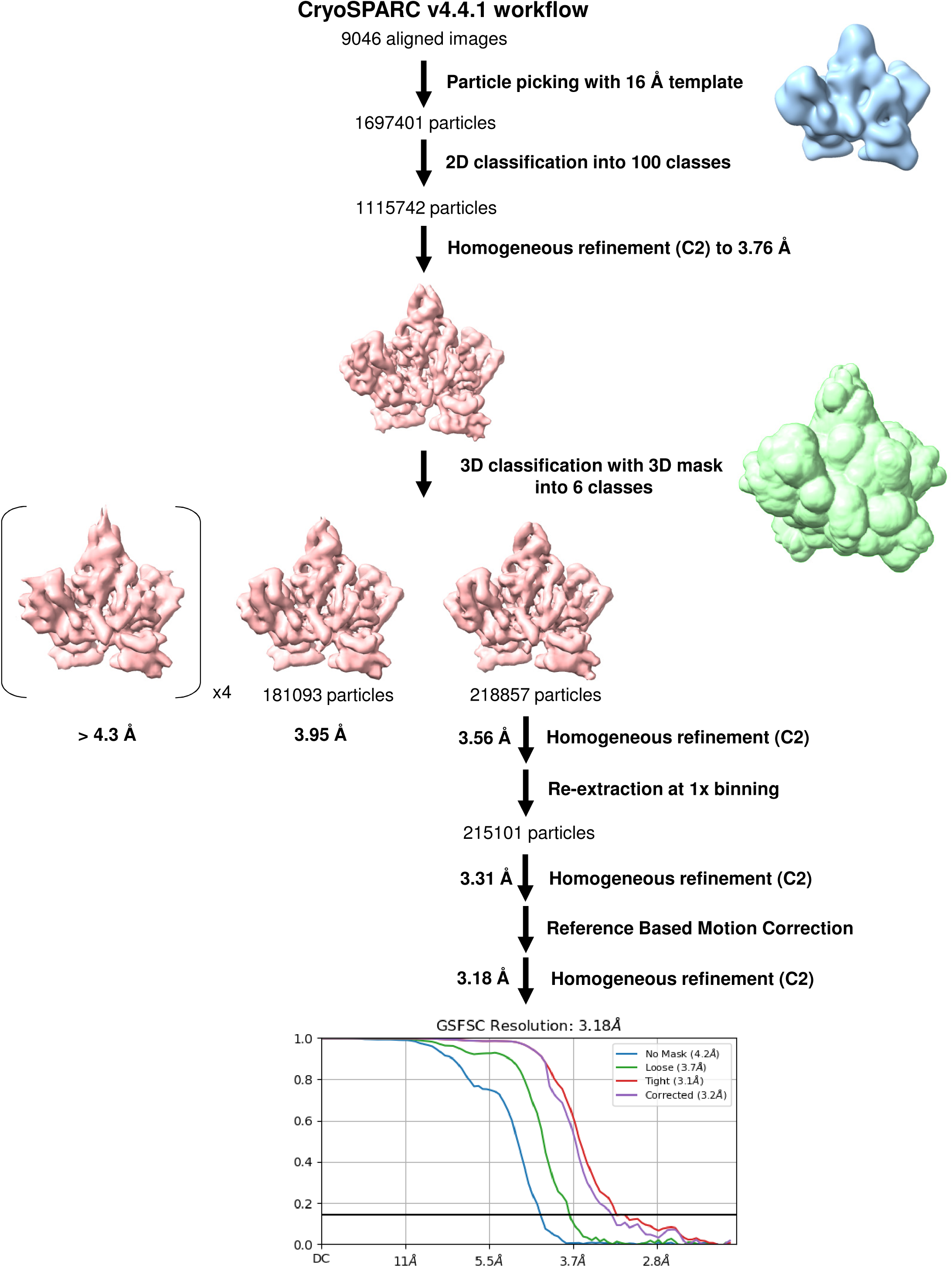
Cryo-EM of GCN2 HRSL domain: workflow for data classification in CryoSPARC; the plot at the bottom shows Fourier shell correlation curves (FSC) as a function of resolution for the 3.18 Å cryo-EM map used for structure determination.

**Figure S2.**
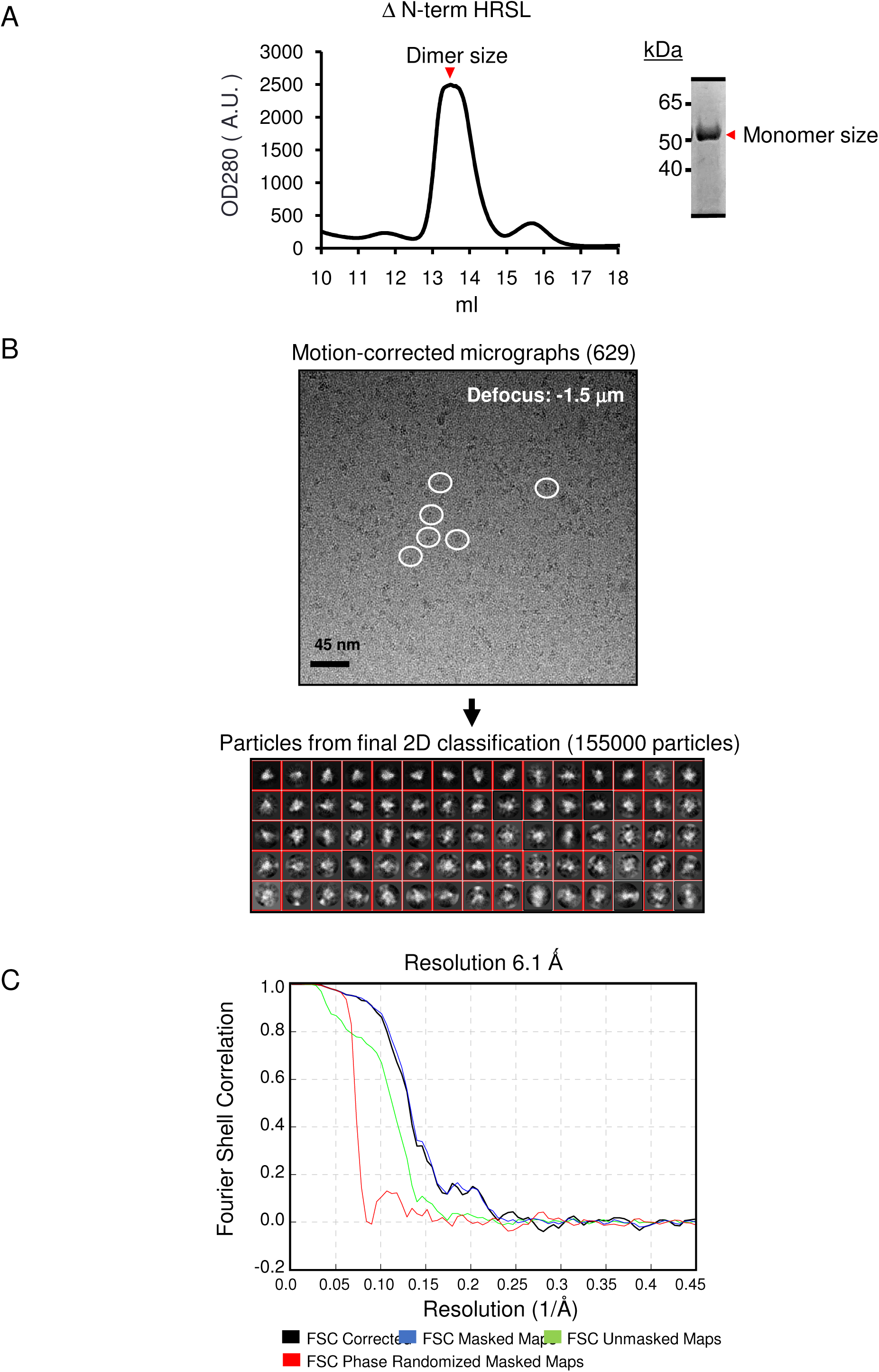
Purification and single particle Cryo-EM analysis of ΔN-term HRSL. A) left-FPLC trace of purified ΔN-term HRSL from Superdex 200 Size Exclusion Chromatography at molecular weight consistent with dimer ∼110 kDa. Right-Coomassie stained SDS-PAGE with purified ΔN-term HRSL collected from indication fraction. Denatured protein runs on gel at expected monomer size ∼55 kDa. B) Top panel: Representative micrograph for ΔN-term HRSL single particle Cryo-EM. White circles-Representative selection of protein particles. ∼155,000 particles were auto-picked and extracted from 629 motion-corrected micrographs and grouped into primary 2D classes and further filtered with additional rounds of 2D classification. Bottom panel: Red boxes from the Final 2D classification yielded ∼46,000 particles and used for 3D initial model. C) Gold-standard Fourier shell correlation (FSC) curve generated from two independent half maps contributing to the 6.1 A resolution of ΔN-term HRSL domain using Relion 3.0.6.

**Figure S3.**
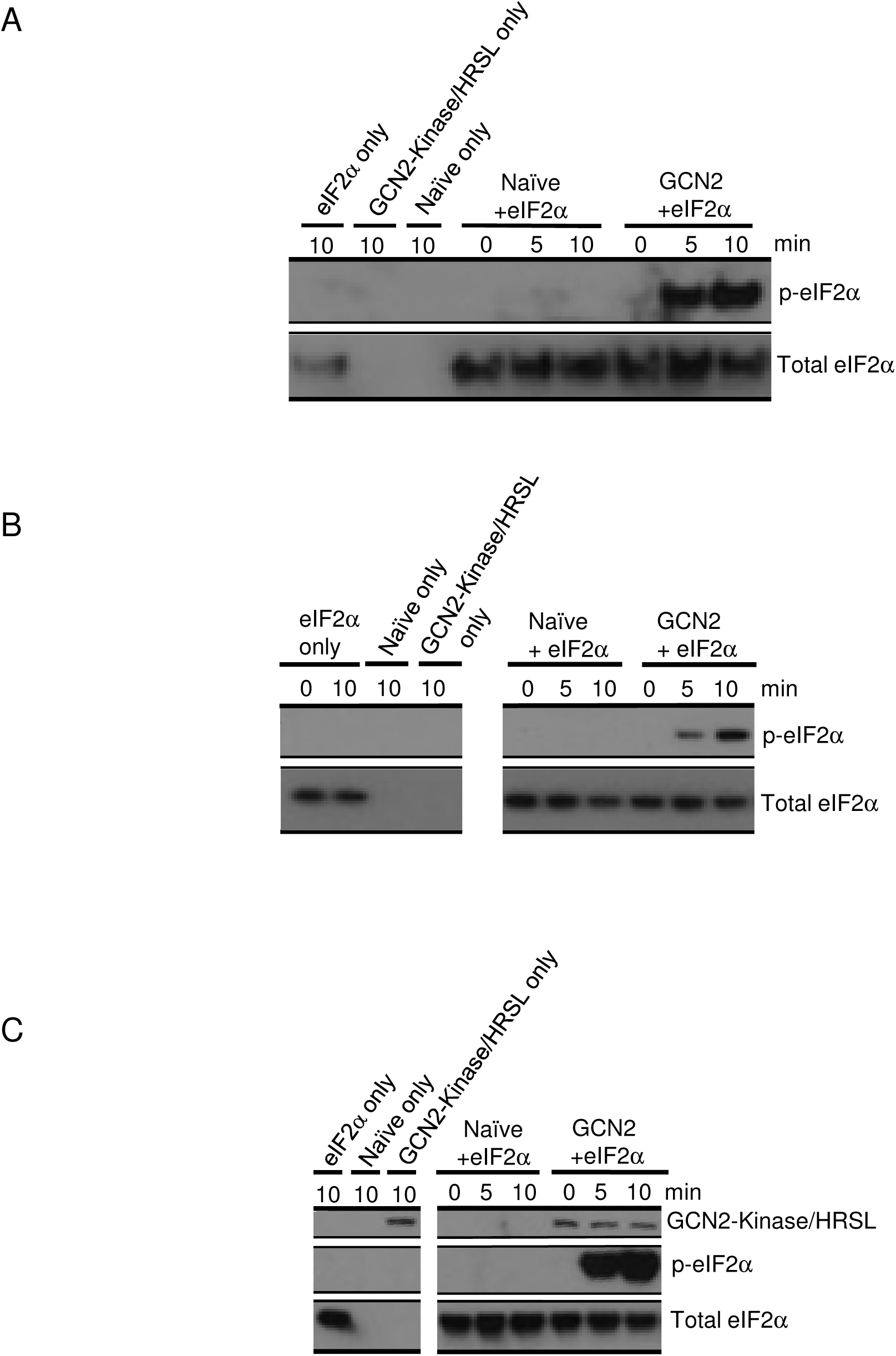
Naïve cell eluate and GCN2 Kinase/HRSL *in vitro* kinase assays. A) Western blot for total eIF2α and phosphorylated eIF2α levels from in vitro kinase assays. eIF2α only lane: kinase assay with only eIF2α substrate present. GCN2 only lane: kinase assay with GCN2 Kinase/HRSL eluted from pulldown only. No eIF2α substrate present. Naïve only lane: cell lysate from naïve cells (cells were not transfected with plasmid bearing GCN2 Kinase/HRSL) underwent a mock pull-down and elution with flag peptide similar to GCN2 Kinase/HRSL pulldown. Performed kinase assay with naïve cell eluate only (No eIF2α or GCN2 Kinase/HRSL eluate present) (B) replicate. (C) Replicate with western blot for GCN2 Kinase/HRSL using anti-flag antibody.

**Figure S4.**
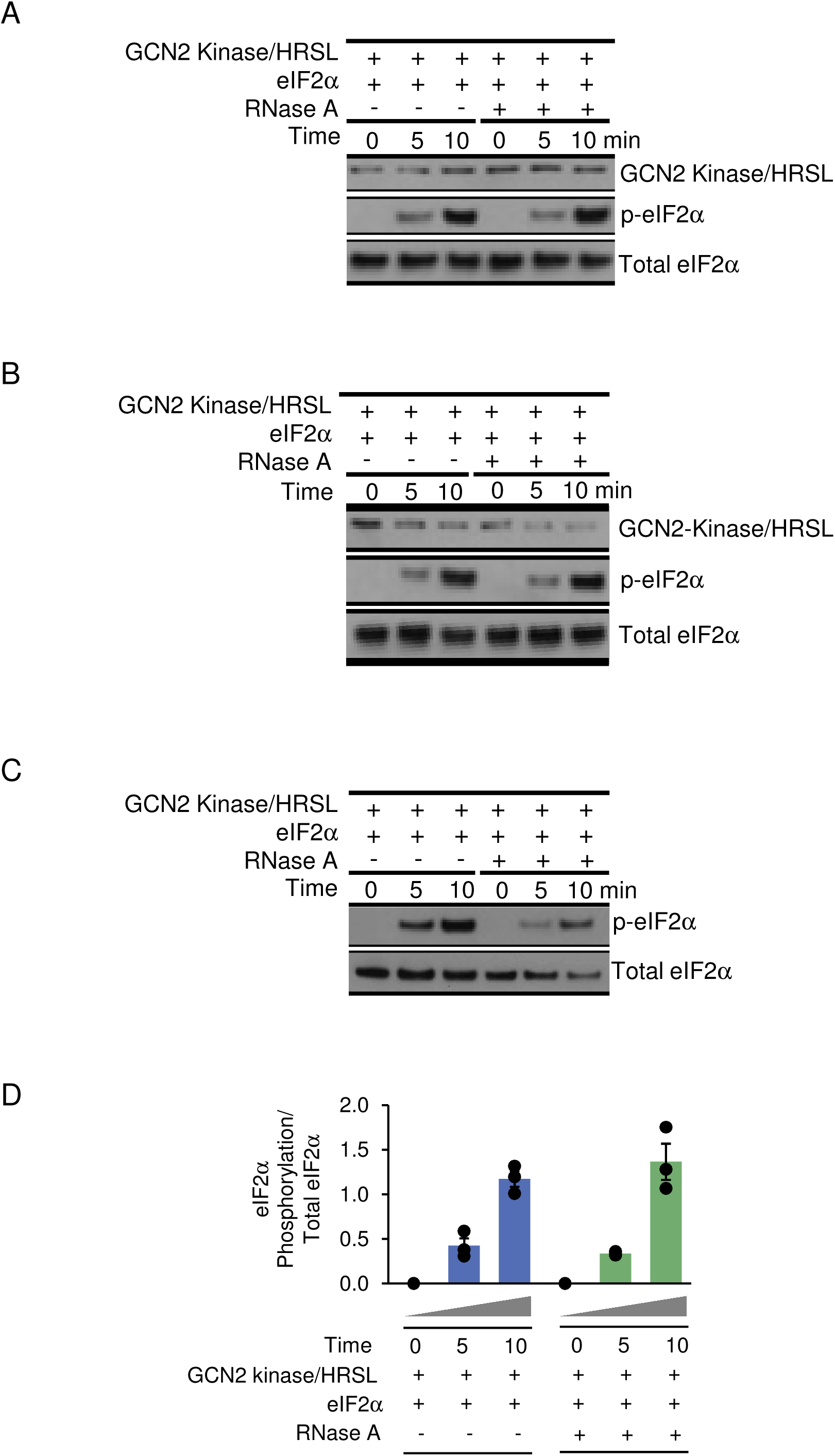
*In vitro* kinase assay with GCN2 Kinase/HRSL interface mutants. **A-B**) Western blots of biological replicates showing dimer interface mutants decrease GCN2 Kinase/HRSL kinase activity related to Fig. 6 A-B.

**Figure S5.**
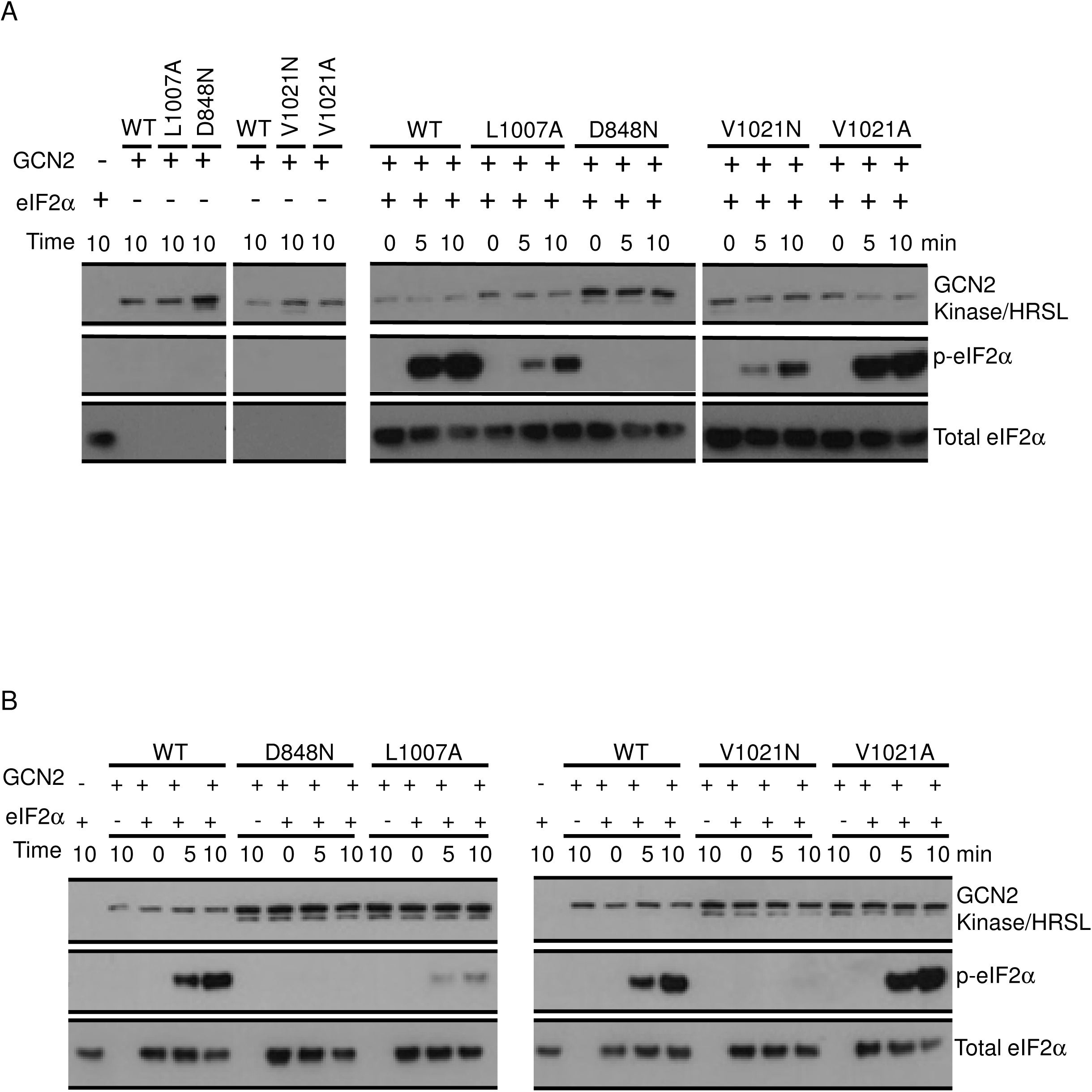
RNase A effect on GCN2 Kinase/HSRL kinase activity. **A)** Western blot for eIF2α phosphorylation from in vitro kinase assay with GCN2 Kinase/HRSL +/- RNase A treatment (350 nM final concentration). Western blot for flag-tagged GCN2 Kinase/HRSL controls for equal loading. **(B-C)** Biological replicates related to **A**. **D)** Results for three replicates (A-C) quantified and normalized to total eIF2α.

## Materials and Methods

### Cloning

GCN2 sequence was amplified from *Kluyveromyces marxianus* genomic DNA, a gift from Prof. Fred Hughson (Princeton University) and inserted into pET11-SUMO3GFP vector, also a gift from Prof. Fred Hughson. Residues 996-1511 of *Kluyveromyces marxianus* GCN2 (kmGCN2) were amplified as GCN2 HRSL construct. Residues 578-1511 of kmGCN2 were amplified as GCN2 Kinase/HRSL construct. Deletions and point mutants (Table S3) were generated using one primer site-directed mutagenesis. All constructs used in this study were verified by DNA sequencing.

### Overexpression and Protein Purification

pET11-SUMO3GFP vector containing GCN2-HRSL with an amino terminal His_6_- SUMO tag was transformed in *E.coli* BL21 (DE3)-CodonPlus RIPL (Agilent) and grown to an OD600 0.6-1.0 in Luria-Bertani (LB) Medium, at 37 °C and induced with 0.2 mM isopropyl-β-D-thiogalactoside (IPTG) for an expressed overnight at 22 °C in 6x1L of LB. The cells were harvested and resuspended in Lysis buffer containing 500 mM NaCl, 20 mM TRIS**•**HCl (pH 8.0), 10 mM Imidazole, 2 mM MgCl_2_, 5 mM β-mercaptoethanol (BME), 10% (v/v) glycerol, and supplemented with 1x *COMPLETE* protease inhibitor cocktail (Roche). Cells were lysed with an EmulsiFlex C3 (Avestin) while chilled. Crude lysates were clarified by high-speed centrifugation at 35,000 g for 30 min at 4 °C. Clarified lysates were loaded onto 3 mL His60 Ni Superflow Resin (Clonetech). Before use, His60 Ni Superflow Resin was washed with lysis buffer. Once protein bound, resin was rinsed with alternating high salt and low salt washes 3x. High salt wash buffer was same as lysis buffer listed above. Low salt wash buffer contained 150 mM NaCl, 20 mM TRIS**•**HCl (pH 8.0), 10 mM imidazole, 2 mM MgCl_2_, 5 mM BME, and 10% (v/v) Glycerol. Resin and protein were washed once more with Gel Filtration buffer: 350 mM NaCl, 20 mM TRIS**•**HCl (pH 8.0), 2 mM MgCl_2_, 5 mM BME, and 5% (v/v) Glycerol. The His_6_ tag was removed with SUMO-protease (provided by Prof. Fred Hughson, Princeton University). Incubations lasted 3-4 h at 4 °C. Cleaved proteins were collected and purified using Superdex 200 size-exclusion chromatography with Gel Filtration Buffer (using 5 mM DTT instead of β-mercaptoethanol). All proteins were purified to more than 95% purity and protein concentrations were quantified from protein sequence using OD280 absorbance.

### Cryo-EM Sample Preparation

Cryo-samples were prepared in Vitrobot Mark IV (Thermo Fisher), which was set to 4 °C with 100% humidity. Buffer for cryo-EM sample preparation contained 350 mM NaCl, 20 mM TRIS**•**HCl (pH 8.0), 2 mM MgCl_2_, 5 mM DTT, 0.1% Triton-X-100 and less than 1% Glycerol, with final protein concentration of 8 mg/ml. Four μL of GCN2 HRSL solution was applied to the surface of glow-discharged Quantifoil R 1.2/1.3 Cu 300 mesh grids with Holey Carbon. After waiting for 40 s, grids were blotted for 4 s with a blot force of 0 and rapidly plunged into pre-cooled liquid ethane for vitrification.

### Cryo-EM Data Acquisition for ΔN-term HRSL

Six hundred twenty nine movie stacks were automatically collected from 3 individual sessions by EPU on a 300 kV Cs-corrected Titan Krios using a K2 Summit detector (with GIF Bio-Quantum Energy Filters, Gatan). Raw movies were collected in K2 super-resolution mode at a magnification of 105,000x with a pixel size of 1.114 Å. The total exposure time was set to 5.6 s with 0.175 s per frame to generate 32-frame gain-normalized mrc stacks. The total dose for a stack is 40 e−/Å^2^. Defocus values were set from −1 to −2.5 mm.

### Cryo-EM Data Processing for ΔN-term HRSL

Movie stacks were motion corrected using Motioncor2 (1) (UCSF), Contrast transfer function (CTF) information was estimated from non-dose-weighted images using Gctf (2). Particles were picked using RELION 3.0.6 (3). Manually picked particles were grouped into 2D classes to use as a reference for Auto-picking using particle size 120 Å. ∼155,000 particles were auto-picked and extracted using particle box size of 160 pixels. These particles underwent multiple rounds of 2D classification to filter out ice spots, background, or protein aggregates. ∼42,000 particles were left to generate the final 3D reconstruction, which was further refined to generate a 3D model yielding a 6.1 Å map using the gold-standard Fourier shell correlation (GSFSC) with a cut-off at 0.143. Since only the map was needed for our purposes, no further refinement, model building, or validation was required.

### Cryo-EM Data Acquisition for 3.2 Å GCN2 HRSL

9048 raw movies on were collected with EPU version 2.14 on a Titan Krios G4 300 kV TEM equipped with a Falcon 4 direct electron detector in EER format. Data was collected in counted mode, with the magnification of 75,000x and a pixel size of 1.1 Å. The total dose was set is 40 e^−^/Å^2^ with exposure time of 5.9 s. Defocus values were set from −1 to −2.5 μm.

### Cryo-EM Data Processing for 3.2 Å GCN2 HRSL

Data processing was performed using CryoSPARC v4.4.1 (4) (Figure S1). From EER files, 40-frame gain-normalized mrc files were generated at 1.1 Å nominal resolution applying gain reference. Micrographs (summarized in Fig. 1D and Fig. S1) were motion corrected using Patch Motion Correction, and contrast transfer function information was estimated with Patch CTF. Micrographs contained visible particles, whose size and shape resembles the AlphaFold (5) dimer of GCN2-HRSL (Fig. 1D). A fraction of the dataset was first used for pre-processing and assessing HRSL macromolecular shape and homogeneity. Blob picking was performed from the first 906 micrographs with the minimal size of 50 Å and maximal size of 120 Å, yielding 2,066,984 particles. Inspect picks was run selecting particles with NCC larger than 0.520 and Local power parameters ranging from 80 to 404 totaling in 425,793 particles. During particle extraction as 2x binned stack (nominal resolution 2.2 Å/px), the particles were excluded that were too close to the image edge, resulting in a 384,902-particle stack with the box size of 100 pixels. This stack of particles was 2D classified into 100 classes with window inner radius 0.75, number of online-EM iterations 80 and number of full iterations 4. Ten best 2D classes were selected based on resolution, ECA and high-resolution features resulting in 63,879 particles. Star file for this stack was generated with script Csparc2star from Pyem package (6) and used to generate *ab initio* model in RELION 4.0 (3) that was subsequently refined without masking to 5.7 Å resolution. Resulting map low-pass filtered to 16 Å was very similar to the 16 Å map generated from the HRSL AlphaFold model (Chimerax correlation between the two is 0.986); the latter was used for expedited template picking from the full dataset of 9048 micrographs.

To process the complete dataset, a reference map was calculated from the AlphaFold model of HRSL and 50 equally spaced 2D projections were generated. For low-resolution template-based particle picking, 2D templates and micrographs were low-pass-filtered to 16-Å resolution. Template particle picking with the particle diameter set to 120 Å and the minimal separation distance set to 90 Å yielded 9711124 particles. We selected particles with NCC score parameters larger than 0.500 and Local power parameters ranging from 341 to 606, amounting to 1,697,401 particles. During particle extraction, the particles were excluded that were too close to the image edge, resulting in a 1,585,862-particle stack with the box size of 160 pixels. This set of particles was subjected to 2D classification into one hundred classes with the maximum alignment resolution set to 5 Å, batch size per class set to 500 per iteration, number of online iterations set to 100 and number of online full iterations to 5.

After discarding particles that belong to low-resolution or non-particle classes, 1,115,742 particles from 54 classes (Fig. 1D) were selected for a round of Homogeneous refinement, which yielded a 3.76 Å three-dimensional map. Refinement with C2 symmetry yielded a better-quality map than that without symmetry. Each step in subsequent analyses was run applying C1 or C2 symmetry group, and the results were compared. Maps with the C2 symmetry were superior in map connectivity to the ones obtained without symmetry at all steps.

To resolve conformational heterogeneity and improve the resolution of the map(s), 3D variability analysis and 3D classification were performed. The 3D variability analysis has not revealed a continuous family of 3D structures. Particles, solvent mask and alignment values from Homogeneous refinement were used as an input to 3D classification into six classes with the high-resolution limit of 6 Å. The highest-resolution 3D class (3.56 Å, 218,857 particles) had the most well resolved and continuous secondary structure, including the N-terminal 4-helix bundle. Refined coordinates and other parameters were re-extracted into a 215,101-particle stack with the box size of 200 pixels (particles close to the box edge were excluded). Homogeneous refinement yielded a 3.31 Å map, which was subjected to the Reference-Based Motion Correction. After the correction and Homogeneous refinement, the final map had the gold-standard Fourier shell correlation resolution of 3.18 Å. We noticed a size discrepancy between the 1.1 Å/px map and the AlphaFold model. To correct the pixel size, we generated maps with pixel sizes ranging from 1.00 Å to 1.20 Å with the step of 0.01 Å using EMAN2 package (7). Using the GCN2-HRSL AlphaFold structure as a starting model, 5 cycles of rigid-body refinement, morphing and minimization were performed against each map using phenix.real_space_refine v1.20rc3 (8). The best fit (CC_mask) and stereochemistry statistics were obtained for the map with the pixel size of 1.04 Å, and the corresponding map was used for the following steps of structure refinement.

### 3.2 Å GCN2 HRSL Model Building and Refinement

Model building was done manually in Coot using AlphaFold dimer prediction as template. Refinement was done with Cryo-EM Real Space Refinement in Phenix (8) with minimization global, rigid body, local grid search, and ADP. Max iterations of 100, 5 macro cycles, target bonds RMSD of 0.01, and target angles RMSD of 1.0. We used secondary structure restraints and NCS. Phenix Cryo-EM Comprehensive Validation statistics are listed in Table S1.

### AlphaFold prediction for GCN2 HRSL dimer

Conducted using locally installed AlphaFold-Multimer v2.1.0 (9). Starting model of the homodimer of the C-terminal domain (a.a. 970-1510) was obtained for GCN2 HRSL. First five models were obtained for a monomer, the highest per-residue confidence metric called “predicted local-distance difference test score” (pLDDTs) was 87.4, indicating a high likelihood of agreement with an experimental structure (5). For the set of five predicted dimers, a different metric was reported, the weighted combination of model accuracy and interfaces accuracy (pTM+ipTM) score. The highest of five pTM+ipTM values was 0.88, indicating the high likelihood of the accurate prediction of intradimer contacts (9).

### Electrostatic Conservational Analysis

Sequence conservation and 3D electrostatic conservation mapping was conducted using SEQMOL-Kd software (BiochemLabSolutions.com). Multiple sequence alignments were first generated using built-in MUSCLE 3.7 (10) using 193 unique sequences of GCN2, from yeasts to human, and 859 unique sequences of tRNA synthetase His, also from yeasts to human. The resulting sequence alignments were used to assign per-amino acid charge conservation, and for computation of conservation-aided electrostatic potentials. Final PDB files were prepared in SEQMOL-Kd and visualized in UCSF Chimera (11).

### GCN2 overexpression in mammalian cells and immunoprecipitation

pcDNA4.TO vectors containing kmGCN2 Kinase/HRSL (WT or mutants) were transfected into HEK293T cells. Cells were a gift from Yibin Kang Laboratory (Princeton University). HEK293T cells were transfected at 60-70% confluency. 30 μg of plasmid was added to 3 ml of opti-MEM and allowed to sit at RT for 5 min. Simultaneously, 40 μl of Lipofectamine 2000 was added to 3 ml of opti-MEM and allowed to sit at room temperature for 5 min. The two mixtures were mixed and allowed to sit at room temperature for 15 min. The total 6 ml of opti-MEM mixture was added to 24 ml of fresh DMEM (Gibco) + 10 FBS for a final volume of 30 ml. Media was not replaced after transfection. Cells were grown at 37 °C with 5% CO_2._

Cells were harvested 18 hrs post transfection by scraping with cold PBS. Cells were spun down at 500 g for 5 minutes and pellets were resuspended in 500 μl of Lysis Buffer (350 mM NaCl, 20 mM HEPES pH 7.5, 1 mM DTT, 5% glycerol, 1x protease inhibitor cocktail (Roche)). Cells were lysed via sonication with 9 pulses twice, on ice. Lysate was spun at 15,000 g for 10 min before adding to pre-equilibrated Anti-DYKDDDDK G1 Affinity Resin (Genscript). For each IP, 100 μL of resin slurry was washed with 1 ml of lysis buffer and spun down at 500 g for 30 s. This step was repeated 3x. Samples were resuspended in 300 μl of lysis buffer. 500 μL of clarified lysate was added to the resin and was gently rotated for 30 min at RT. Resin was spun at 500 g for 30 s to remove unbound protein. The resin was washed 3x by re-suspending in 1 ml of was buffer and spun at 500 g for 30 s. The resin was then resuspended in 500 μl Elution Buffer (Lysis buffer with 0.1 mg/ml of DYKDDDDK peptide (Genscript)) Resin was gently rotated for 5 min at RT and spun down at 1000 g for 1 min. Eluate was transferred to 10 kDa concentrator and concentrated to ∼100 μl. This eluate was then immediately used for *in vitro* kinase assays.

### eIF2-α *in vitro* kinase assays

Reactions were performed similar to description of Inglis et al. (12). Reaction buffer contained 100 mM NaCl, 50 mM HEPES (pH 7.4), 5 mM MgCl_2_, 1 mM DTT, and 6.25 mM β-glycerophosphate. GCN2 Kinase/HRSL eluate was added to reaction buffer with 1 mg/ ml BSA. Final concentration of 250 nM eIF2α (LifeSpan Biosciences) was added to the reaction buffer. Activator mix (0.5 mM ATP and 18.75 mM MgCl_2_) was added to the reaction and the tube was then moved from ice to 32 °C water bath. At each time point 10 μl was removed after mixing thoroughly and stopped with 4X SDS+DTT (Sigma). Samples were boiled and analyzed by Western Blot (WB).

### RNase A treatment

Five to ten μl of eluted GCN2 WT Kinase/HRSL were incubated with 350 nM RNase A for 15 min at room temperature (50 mM HEPES pH 7.4, 100 mM NaCl, 5 mM MgCl_2,_ 1 mM DTT, 6.25 mM β-glycerophosphate + 1 mg/ml BSA). Reactions were moved to ice. Final concentration of 250 nM eIF2α (LifeSpan Biosciences) was added to reaction buffer. Activator (Final concentration of 0.5 mM ATP and 18.75 mM MgCl_2_) was added to the reaction and the tube was then moved from ice to 32 °C water bath. During time points of 0, 5 and 10 min, 10 μl was removed after mixing thoroughly and stopped with 4X SDS+DTT stop-denaturing buffer. Samples were boiled and analyzed by WB.

### Western Blotting

Samples were denatured in 4X SDS+DTT, separated by 10% (v/v) BisTris PAGE (NuPAGE), and transferred to PVDF membranes (Life Technologies). Following 15 minutes incubation with blocking buffer [5% (w/v) nonfat dry milk in TRIS-Buffered Saline and Tween-20 (standard TBST)], the membranes were probed with the following primary antibodies: 1) mouse anti-human total eIF2α (Cell Signaling Technology) at 1:1,000, 2) rabbit anti-human eIF2α-phosphoS51 (Abcam) at 1:1,000 3) mouse anti-Flag M2 (Sigma) at 1:2,000. Antibody treatments were followed by horseradish peroxidase-conjugated anti-rabbit or anti-mouse secondary antibodies (1:5,000 or 1:10,000 respectively, Jackson ImmunoResearch) (Table S3). The membranes were washed, blots were developed with enhanced chemiluminescence Western blotting detection reagents (GE Healthcare Life Sciences), and gels were exposed to X-ray film for visualization.

### Error analysis

Error bars in solution assays represent S.E. from three independent experiments. P-values were calculated using Welch’s two-tailed unpaired t-test as described in our prior work (13).

---

**Table S1.** Cryo-EM Data Collection and Refinement Statistics.

| | *K. marxianus* GCN2-HRSL | ΔN-term GCN2 HRSL |
| --- | --- | --- |
| **Data collection** |  |  |
| Grid Type | QuantiFoil R 1.2/1.3 Cu<br>300 mesh<br>Holey Carbon Films | QuantiFoil R 1.2/1.3 Cu<br>300 mesh<br>Holey Carbon Films |
| Microscope/Detector | Titan Krios / G4 | Titan Krios / Gatan K2 |
| Voltage | 300 kV | 300 kV |
| Magnification | 75,000x | 105,000x |
| Recording mode | Counting Mode | Super Resolution Counting Mode |
| Defocus | -1.0 – -2.5 μm | -1.0 – -2.5 μm |
| Pixel size | 1.04 Å | 1.04 Å |
| Total dose | 40 e<sup>-</sup>/Å<sup>2</sup> | 40 e<sup>-</sup>/Å<sup>2</sup> |
| Number of frames/movie | 40 | 32 |
| Total exposure time | 5.96 s | 5.6 s |
| Number of micrographs | 9046 | 629 |
| Number of particles picked | 1,585,862 | 155,000 |
| Number of particles after 2D | 1,115,742 | 46, 000 |
| Particles used for final map | 215,101 | 46, 000 |
| Map resolution (GSFSC 0.143) | 3.18 Å | 6.1 Å |
| PDB | 9BF3 | - |
| EMDB | 44489 | - |
| **Real-Space Refinement** |  |  |
| Resolution (Å) | 3.2 | - |
| Map CC_mask | 0.86 | - |
| *B* factor (Å<sup>2</sup>) for Protein | 119 | - |
| **Validation** |  |  |
| All-atom class score | 3.66 | - |
| Rotamer outliers (%) | 0.00 | - |
| CaBLAM outlier (%) | 0.21 | - |
| MolProbity score | 1.18 | - |
| **Ramachandran plot** |  |  |
| Favored (%) | 97.92 |  |
| Allowed (%) | 2.08 |  |
| Outliers (%) | 0 |  |
| **RMS deviation** |  |  |
| Bond length | 0.002 |  |
| Bond Angle | 0.417 |  |

**Table S2.**
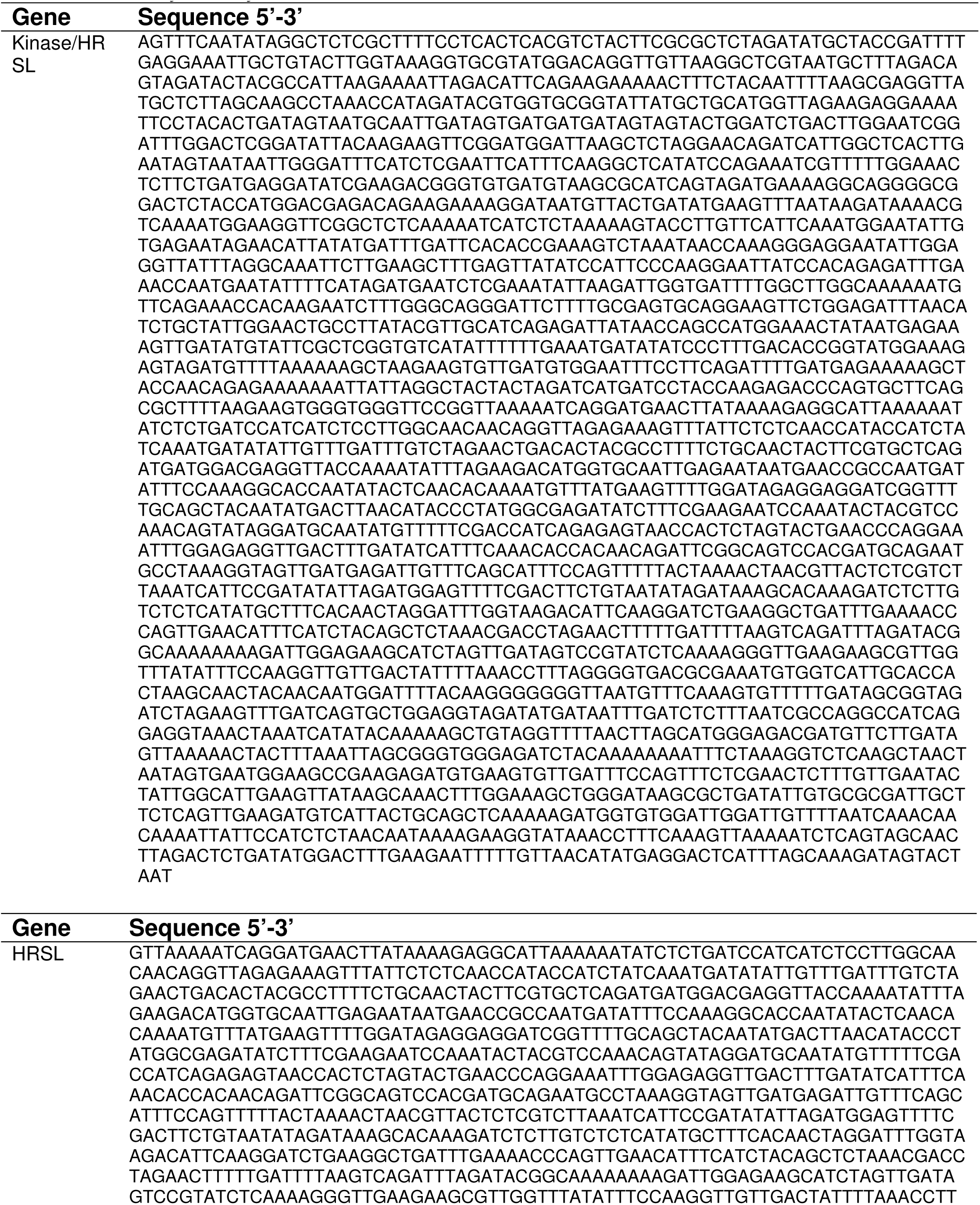
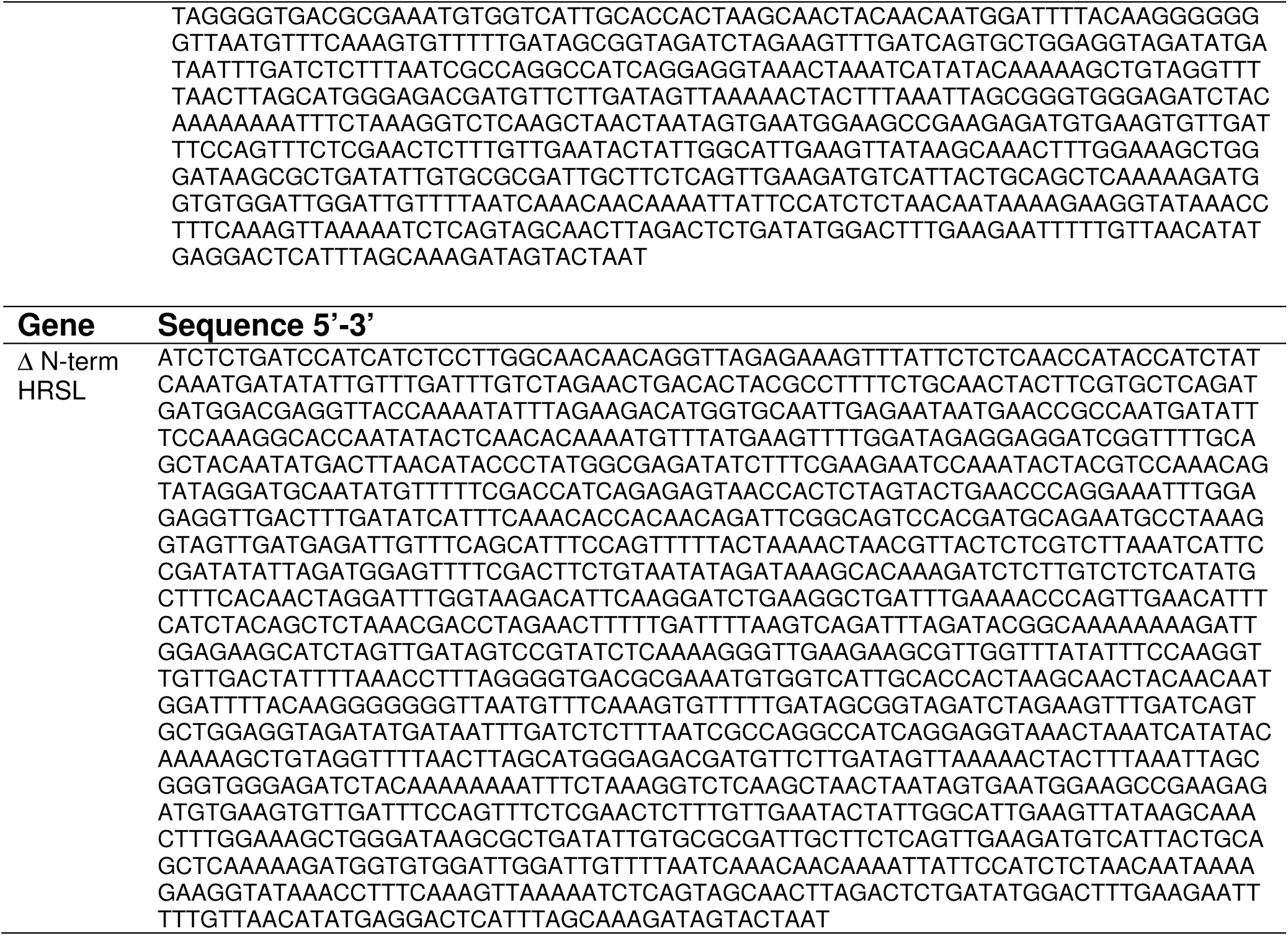
DNA sequences used for recombinant protein expression of *Kluyveromyces marxianus* GCN2.

**Table S3.**
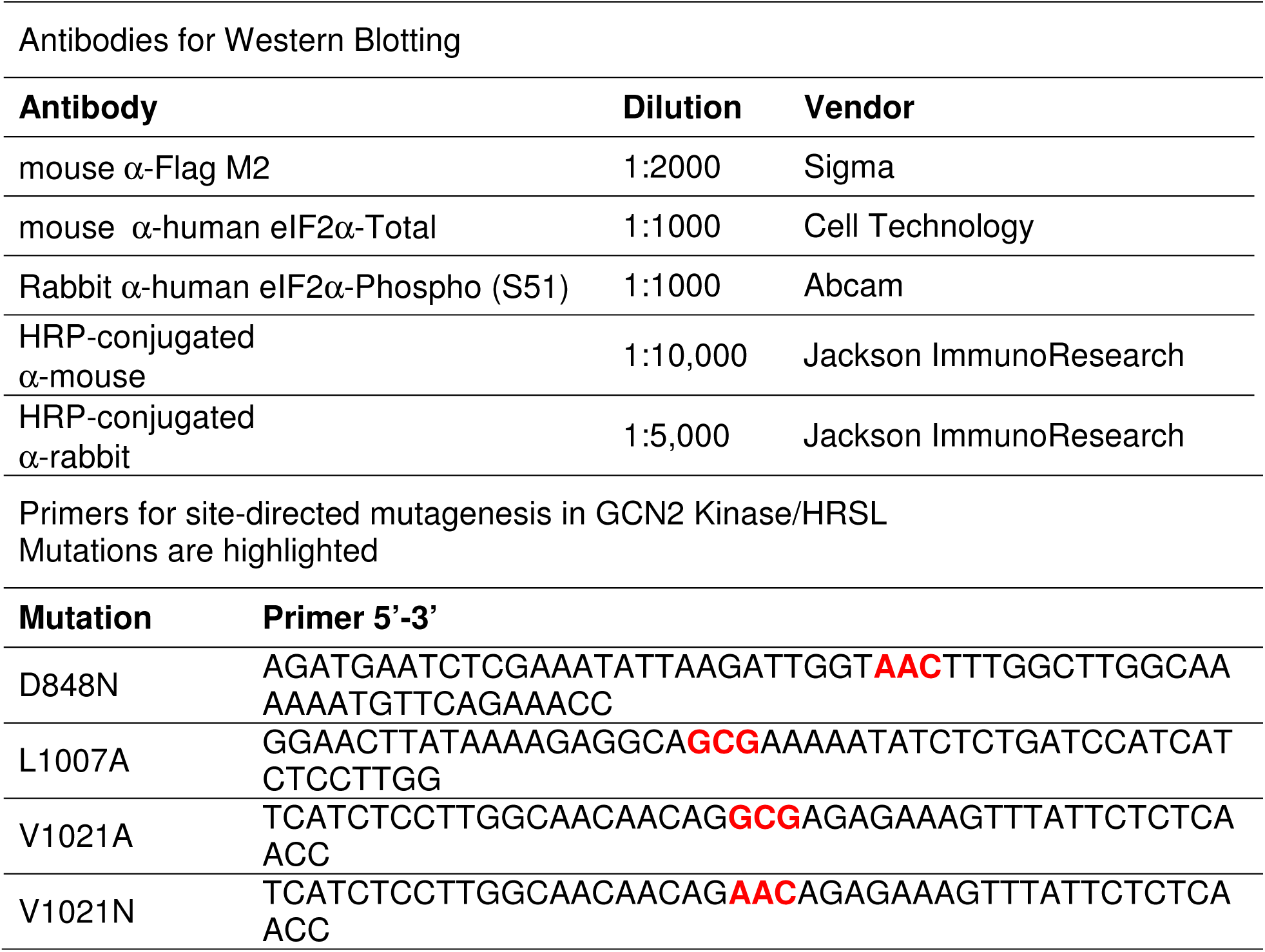
Reagents.

